# Compound screening in primary human airway basal cells identifies Wnt pathway activators as potential pro-regenerative therapies

**DOI:** 10.1101/2024.08.13.606573

**Authors:** Yuki Ishii, Jessica C. Orr, Marie-Belle El Mdawar, Denise R. Bairros de Pilger, David R. Pearce, Kyren A. Lazarus, Rebecca E. Graham, Marko Z. Nikolić, Robin Ketteler, Neil O. Carragher, Sam M. Janes, Robert E. Hynds

**Author notes:** **Correspondence:** Robert E. Hynds Sam M. Janes. These authors contributed equally (Yuki Ishii, Jessica C. Orr).

## Abstract

Regeneration of the airway epithelium restores barrier function and mucociliary clearance following lung injury and infection. Basal cells are tissue-resident airway stem cells that enact regeneration, yet the mechanisms regulating their proliferation and differentiation remain incompletely understood. To identify compounds that promote primary human airway basal cell proliferation, we performed phenotype-based compound screening of 1,429 compounds (from the ENZO and Prestwick Chemical libraries) in 384-well format using primary cells transduced with lentiviral luciferase. 16 pro-proliferative compounds validated in independent donor cell cultures, with several hit compounds activating the Wnt signalling pathway. The effects of compounds on proliferation were further explored in concentration-response, colony formation and 3D organoid assays. Structurally and functionally-related compounds that more potently induced both Wnt activation and basal cell proliferation were investigated. One such compound, 1-azakenpaullone, induced Wnt target gene activation and basal cell proliferation in mice in the absence of tracheal injury. Our results demonstrate the pro-proliferative effect of small-molecule Wnt activators on airway basal cells. These findings contribute to the rationale to develop novel approaches to modulate Wnt signalling during airway epithelial repair.

**Summary statement:** Ishii, Orr and colleagues perform a high-throughput screen of 1,429 compounds in primary human airway epithelial cells, identifying Wnt activating compounds as promoters of proliferation.

## INTRODUCTION

Human airways are lined by a pseudostratified epithelium that acts as a physical barrier and enacts mucociliary clearance to remove inhaled particles and pathogens from the lungs (Bustamante-Marin and Ostrowski, 2017). Maintaining these functions during homeostasis and following injury is essential for lung health. Tissue-resident basal stem cells can self-renew and differentiate to replenish airway epithelial cell lineages (Basil et al., 2020; Rock et al., 2009).

Airway epithelial repair is a feature of a range of respiratory conditions. Influenza infection leads to tracheobronchitis and cytonecrosis, with desquamation of epithelial cells into the luminal space (Taubenberger and Morens, 2008; Walsh et al., 1961). Sloughing of the airway epithelium is also a feature of respiratory syncytial virus (RSV) (Johnson et al., 2007) or rhinovirus infections (Turner et al., 1982). In chronic lung disease, basal cell stress and loss of progenitor function have been proposed as mechanisms underlying chronic obstructive pulmonary disease (Ghosh et al., 2018; Staudt et al., 2014; Wijk et al., 2021). Additionally, basal cells from asthma patients exhibit functional defects *in vitro* (Kicic et al., 2010; Stevens et al., 2008). Dysregulated airway repair can also lead to chronic airway inflammation and fibrosis in epidermolysis bullosa (Lau et al., 2024), and bronchiolitis obliterans-like pathologies in experimental models (O’Koren et al., 2013). Promoting airway epithelial regeneration is also a key aim of future lung regenerative medicine approaches, including tissue transplantation approaches where promotion of the endogenous epithelial infiltration of the graft is desirable, and following airway epithelial cell transplantation. As such, understanding the mechanisms regulating basal cell function and developing approaches to modulate basal cell behaviour are of significant interest (Hynds, 2022).

Drug development is a costly process, with many candidate drugs failing due to a lack of efficacy or toxicity concerns in on- and off-target effects (Sun et al., 2022). Since it takes, on average, 10-15 years before a new medicine can reach patients (Berdigaliyev and Aljofan, 2020), drug repurposing (i.e. identifying new uses for approved medications outside the scope of the original indication) is an attractive approach (Ashburn and Thor, 2004). Molecules that have been found to be safe in preclinical models and clinical trials in another setting increases the chance of success and can reduce the time and investment required to take therapies from the bench to the bedside (Pushpakom et al., 2019).

Airway regeneration can be modelled in primary cell cultures. Human bronchial epithelial cells (HBECs) proliferate in both 2D or 3D culture systems (Hiemstra et al., 2018; Nizamoglu et al., 2023; Orr and Hynds, 2021), and can be induced to differentiate in air-liquid interface (ALI) (Fulcher et al., 2005) or organoid cultures (Danahay et al., 2015). In recent years, the adaptation of cell culture models to 96-, 384- or 1536- format plates have allowed for higher throughput phenotypic screening (Bloom, 2021; Mayr and Bojanic, 2009). Here, we combine primary airway basal cell culture with compound screening to identify novel modulators of airway basal cell proliferation. Compounds that were identified as promoting basal cell proliferation were further validated in both 2D and 3D cell culture models to assess their effects on basal cell proliferation and differentiation.

## RESULTS AND DISCUSSION

### Compound screening in primary airway basal cells

Primary human bronchial epithelial cell cultures were established from endobronchial biopsy specimens. Cells underwent lentiviral transduction with a pHIV-Luc-zsGreen construct and fluorescence activated cell sorting to purify zsGreen-positive cells (**Fig. S1A-C**). This enabled 384-well format screening for modulators of basal cell proliferation by monitoring bioluminescence following the addition of luciferin to the culture medium (**Fig. 1A**). In validation experiments, bioluminescence signals correlated strongly with Hoescht 33342 nuclei counts (**Fig. 1B** and **Fig. S1D**) suggesting that our screening approach successfully reports cell proliferation. As a positive control, we tested Y-27632, a small-molecule inhibitor that is known to induce basal cell proliferation (Horani et al., 2013; Reynolds et al., 2016). As expected, the bioluminescence signal increased following the addition of Y-27632 (**Fig. 1C**). Calculation of the signal window (10.09) and Z’ factor (0.7) between control and Y-27632- treated wells indicated a robust assay window (signal window ≥ 2, Z’ factor ≥ 0.4; Iversen et al., 2012). The hit threshold was set at 1.96 standard deviations above the plate mean (i.e. a mean Z score above 1.96).

**Figure 1:**
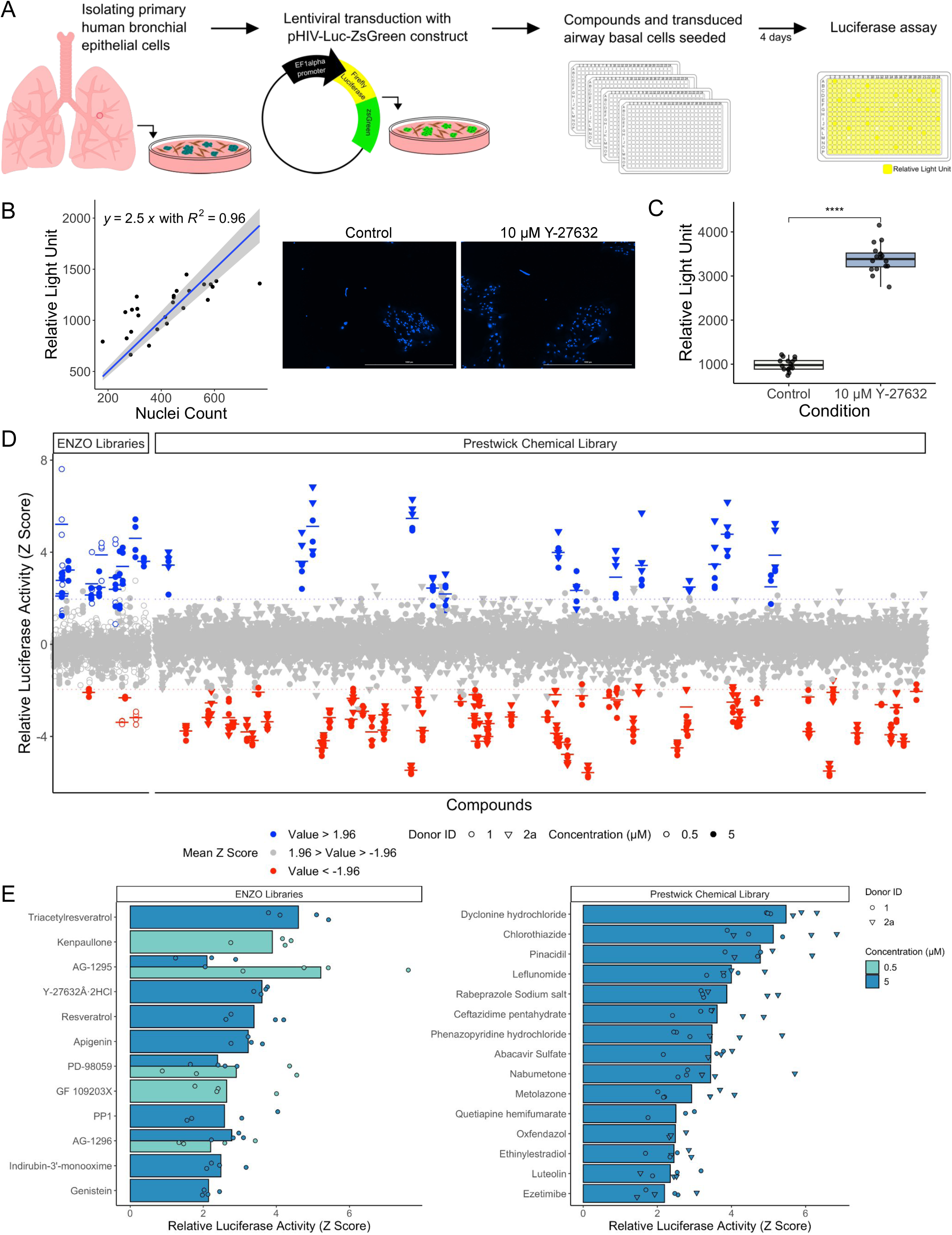
High-throughput screening of 1,429 compounds identifies compounds that modulate airway basal cell proliferation. A) Schematic representation of the 384-well format screening approach. Primary human airway basal cells were transduced with a pHIV-Luc-zsGreen lentivirus to enable luciferase monitoring of cell proliferation. Cells were seeded in 384-well format and treated with compounds from the ENZO chemical library and the Prestwick Chemical library for four days before luciferin was added to the culture medium and bioluminescence was measured. B) Correlation between bioluminescence and nuclei count (left; n = 16 wells, donor ID 1, see also Supplemental Fig. 1D). Representative images of Hoechst 33342 staining of cells following the luciferase assay (right). Scale bars = 1,000 μm. C) Bioluminescence readings in the 384-well format assay comparing control or 10 μM Y-27632 treated wells (n = 16 wells, donor ID 1, Wilcoxon test, ****=p<0.0001). D) 1429 compounds from either an epigenetic modulator, protease and kinase inhibitor library (ENZO Life Sciences) or the Prestwick Chemical library were screened in transduced airway basal cells (n = 2 donors [ID 1 = circles, ID 2 = triangles]) in triplicate. Z Scores were calculated from the relative light unit values. Hit compounds (blue) were identified as those with a mean Z Score above 1.96 standard deviations of the plate average (above the grey dotted line). Red colour indicates compounds with a mean Z Score below 1.96 standard deviations of the plate average. E) Hit compounds identified from screening of the ENZO chemical library (left) and the Prestwick Chemical library (right) [donor ID1 = circles, ID2a = triangles].

We screened a total of 1,429 compounds from the ENZO chemical library (of epigenetic modulator and proteinase and kinase inhibitors) and the Prestwick Chemical library (of compounds that are approved for clinical use by the FDA, EMA and other agencies and have been selected to represent broad pharmacological diversity). We identified 27 compounds (**Table S1**) that increased luciferase activity above the threshold value (**Fig. 1D/1E**), including Y-27632. An additional 61 compounds significantly reduced the number of basal cells (**Fig. 1D**), likely including compounds that inhibit essential processes, inhibit the cell cycle or are toxic to basal cells. Z scores for all compounds tested can be found in **Table S2**. The establishment of a robust airway basal cell proliferation screening protocol further demonstrates the ability to perform screening in primary airway epithelial cells. Our approach adds to previous 96-well format screens for modulators of ciliogenesis in mouse organoids from transgenic mice (Tadokoro et al., 2014), readthrough agents in primary ciliary dyskinesia air-liquid interface cultures (Lee et al., 2021) and anti-viral compounds in healthy air-liquid interface cultures (Sen et al., 2024), as well as a 384-well plate screen for inhibitors of thymic stromal lymphopoietin (TSLP) production performed in commercially available airway basal cells (Orellana et al., 2018).

To validate that hit compounds accelerate basal cell proliferation, we re-ordered them from independent suppliers and performed concentration-response proliferation assays at the same timepoint. In primary human bronchial epithelial cell cultures that had undergone lentiviral transduction with pHIV-Luc-zsGreen (n = 3 donors), 17 hit compounds (10 from the ENZO chemical library and 7 from the Prestwick Chemical library) increased luciferase activity, including Y-27632 (**Fig. 2A, Fig. S2A**). Five of six hit compounds tested from the Prestwick Chemical library in organoids (or ‘bronchospheres’) led to an increase in organoid size (**Fig. S2B**).

**Figure 2:**
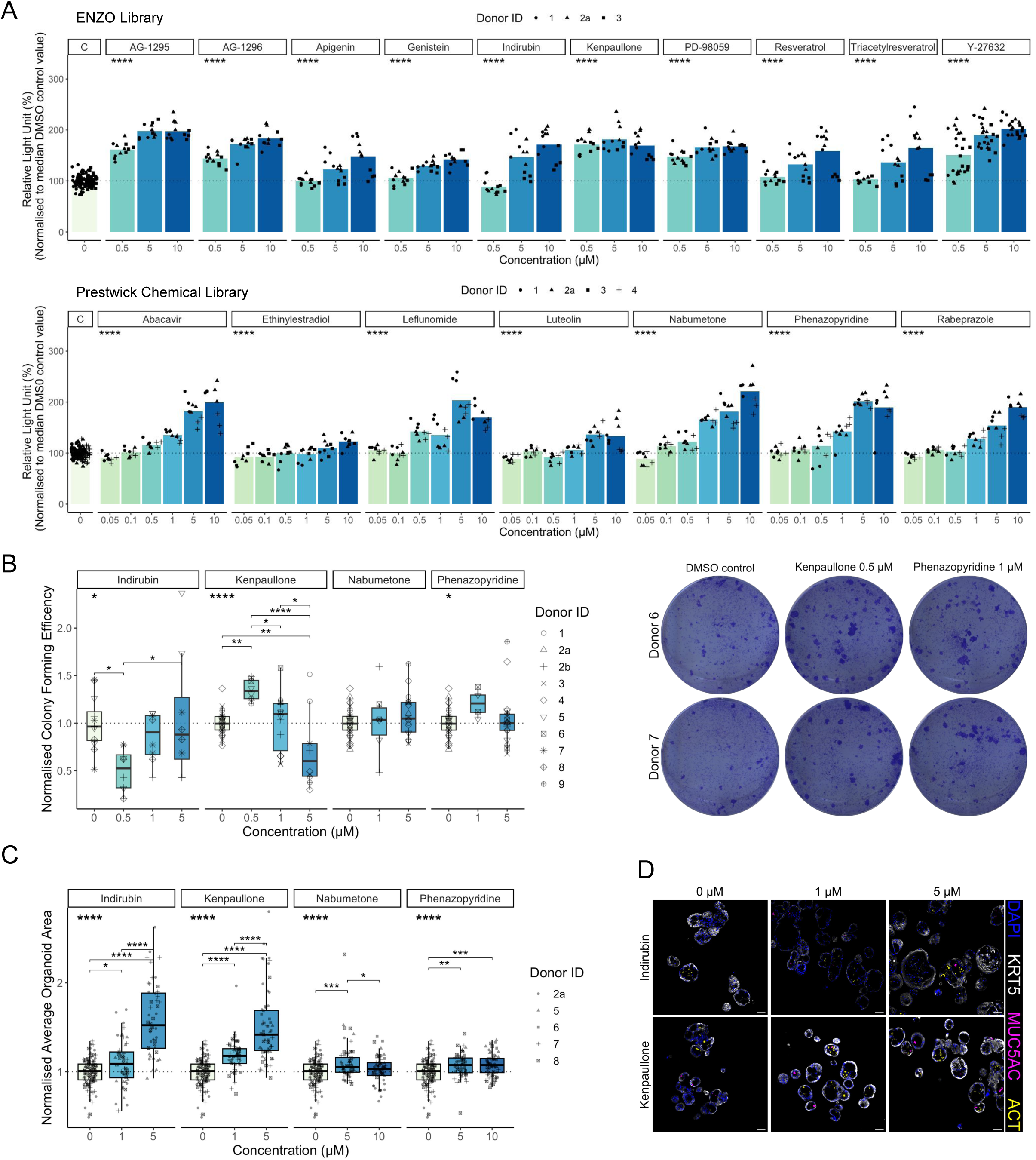
Validation of pro-proliferative compounds in 2D and 3D primary cell culture models. A) Four-day concentration-response proliferation assays in primary human airway basal cells transduced with the pHIV-Luc-zsGreen construct for compounds identified within the ENZO chemical library (upper panel) or the Prestwick Chemical library (lower panel; n = 3 donors per compound; donor ID1 = circles, ID2a = triangles, ID3 = squares, ID4 = cross). An ANOVA was performed per compound, **** = p<0.0001. B) Quantification of airway basal cell colony formation efficiency for four validated hit compounds identified as Wnt activating molecules (left; n=3-9 donors per compound) and representative well images from the 0.5 µM kenpaullone - and 1 µM phenazopyridine-treated wells (right). An ANOVA was performed per compound and significant Tukey’s HSD values are shown. * = p<0.05, ** = p<0.01, **** = p<0.0001. C) Quantification of mean organoid size per well with 12 replicate wells per condition were normalised to mean control well organoid size for each donor (n = 4/5 donors per compound; donor ID1 = circles, ID2a = triangles, ID3 = squares, ID4 = cross). An ANOVA was performed per compound and significant Tukey’s HSD values are shown. * = p<0.05, ** = p<0.01, ***= p<0.001, **** = p<0.0001). Control well data are repeated per compound facet. D) Representative immunofluorescence staining of organoids cultured in the presence of indirubin-3’-monoxime or kenpaullone (n=5, donor ID= 8 shown). Staining for keratin 5 (KRT5; basal cells, white), MUC5AC (mucosecretory cells, magenta) and ACT (multiciliated cells, yellow). Scale bars = 50 μm.

Overall, this screen suggests multiple avenues for future research, including suggesting compounds with re-purposing potential in lung regeneration applications.

### Wnt pathway-activating molecules promote airway basal cell proliferation

Four validated hit compounds - indirubin-3’-monoxime, kenpaullone, nabumetone and phenazopyridine - have been previously reported to activate canonical Wnt signalling. Wnt signalling is involved in the specification of lung progenitor cells (Goss et al., 2009) and the proximal-distal patterning of airways during development (Hashimoto et al., 2012; Shu et al., 2005), and influences post-natal airway epithelial cell proliferation (Aros et al., 2020a; Hsu et al., 2014; Ievlev et al., 2022; Lynch et al., 2016), differentiation (Cooney et al., 2023; Haas et al., 2019; Malleske et al., 2018; Schmid et al., 2017) and the formation of ageing-associated glandular-like epithelial invaginations (Aros et al., 2020b). As such, we further investigated the effects of these compounds in colony formation assays finding that kenpaullone and phenazopyridine increased colony forming efficiency (**Fig. 2B**). We also tested the effects of the ENZO library hit compounds on untransduced primary human bronchial epithelial cells and found that kenpaullone was the compound that most potently induced basal cell proliferation, followed by indirubin-3’-monoxime, apigenin and PD-98059 (**Fig. S3)**. In 3D organoids, the addition of kenpaullone and indirubin-3’-monoxime significantly increased organoid size (**Fig. 2C**). At concentrations of these compounds that increased organoid size, the differentiation potential of basal cells was preserved, with both acetylated tubulin (ACT) and mucin-5AC (MUC5AC) observed by immunofluorescence (**Fig. 2D**), indicative of multiciliated and mucosecretory cell differentiation, respectively.

A previous screening study identified potential lung pro-regenerative therapies by examining the extent to which compounds in the Prestwick Chemical library activated a TCF/LEF reporter in 3T3 mouse embryonic fibroblasts (Costa et al., 2021). Two of the five candidate drugs identified in that study, phenazopyridine and nabumetone, increased basal cell proliferation in both screening (**Fig. 1E**) and validation experiments (**Fig. 2A**). Consistent with our human cell data, the previous study found that phenazopyridine increased the number of mouse lung organoids in cell culture.

Kenpaullone was the most potent pro-proliferative compound identified from the ENZO library screen (**Fig. 2A**) and, of the four reported Wnt activating molecules identified in our screen, induced the highest firefly luciferase activity in HBEC3-KT cells transfected with TOPFlash plasmids (in which firefly luciferase expression is controlled by a minimal promoter and multiple TCF binding sites; **Fig. 3A**). Since more potent and selective derivatives were available for two of the Wnt-activating screening hits identified (kenpaullone and indirubin-3’-monoxime), we compared the effect of kenpaullone to 1-azakenpaullone (Kunick et al., 2004) and 6-bromoindirubin-3-oxime (BIO) (Meijer et al., 2003). Basal cells continued to express the transcription factor TP63, a protein associated with basal cell stem cell potential (Warner et al., 2013), after culture in all three compounds (**Fig. S4A**), and both 1-azakenpaullone and BIO induced higher TOPFlash expression in HBEC3-KT cells than kenpaullone, signifying increased Wnt signalling induction (**Fig. 3B**). This conclusion was further supported by the finding that pretreatment with iCRT14, a small-molecule inhibitor that blocks the transcriptional activity of β-catenin, prevented TOPFlash activation in BIO-treated cells (**Fig. 3B**). We observed higher expression of the Wnt target genes *LEF1* and *AXIN2* in BIO-treated cells (**Fig. 3C**). The expression of *CCND1* and *MYC* were unchanged in kenpaullone, 1-azakenpaullone and BIO treated cells (**Fig. S4B**). All three compounds increased basal cell proliferation as measured by the CellTiter-Glo luciferase assay performed on non-transduced primary airway basal cells (**Fig. 3D**). Moveover, pre-treatment with iCRT14 prevented the increases in proliferation seen with addition of kenpaullone, 1- azakenpaullone or BIO (**Fig. 3E**). Since kenpaullone acts as an inhibitor of both glycogen synthase kinase 3 beta (GSK3β) and cyclin-dependent kinase (Leost et al., 2000; Zaharevitz et al., 1999), this iCRT14 sensitivity suggests that the mechanism of action is canonical Wnt pathway activation. Finally, the compounds increased organoid size in 3D culture in a concentration-dependent manner (**Fig. 3F**). Organoids cultured in medium containing all three compounds displayed both ACT+ ciliated cells and MUC5AC+ mucosecretory cells (**Fig. 3G**) indicating their retained ability to undergo airway differentiation in the presence of these compounds.

**Figure 3:**
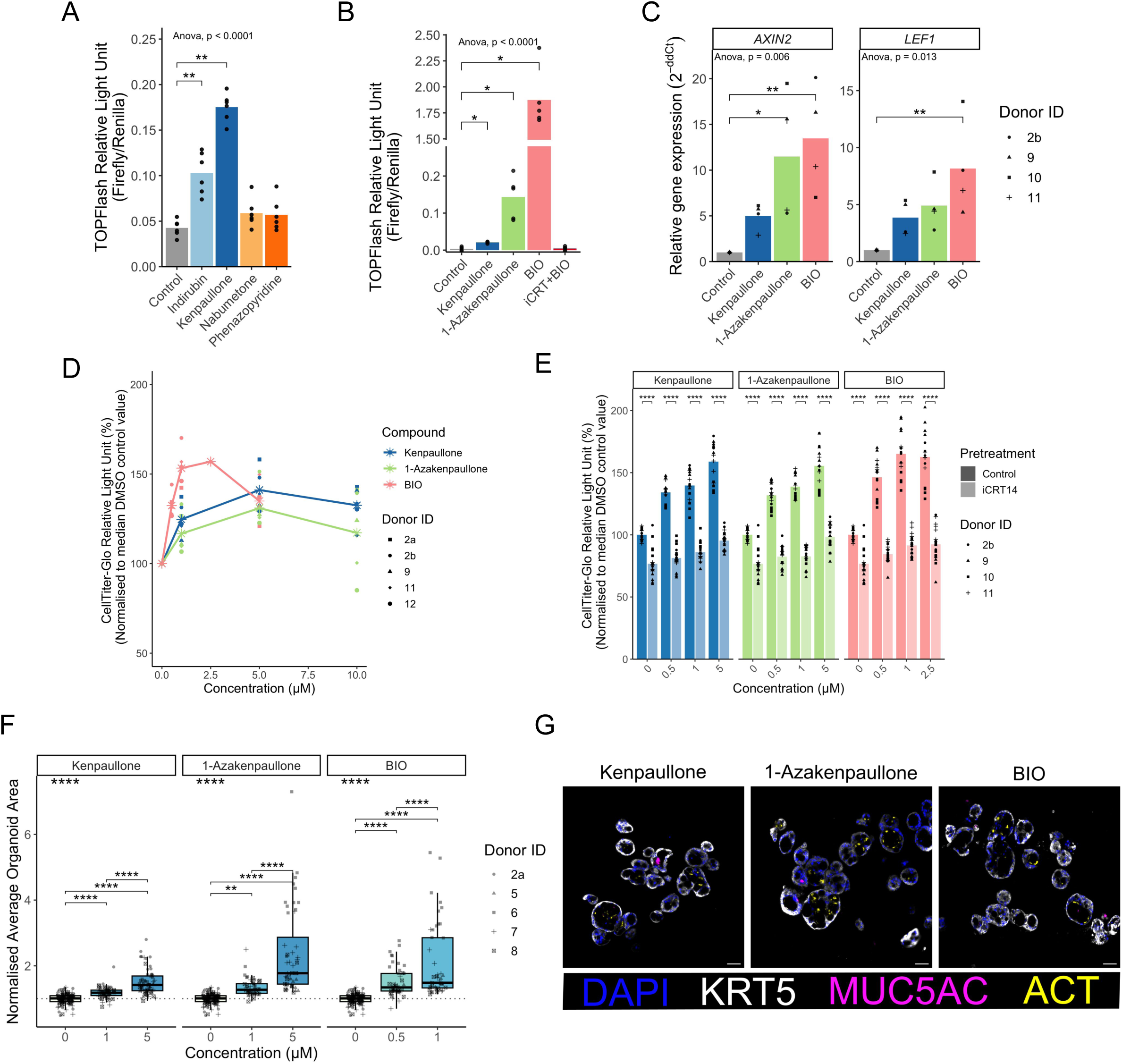
Investigation of Wnt pathway activating compounds as promoters of airway basal cell proliferation. A) Quantification of Wnt activity using the TOPFlash assay. HBEC3-KT cells were transfected with TOPFlash plasmid, re-plated in a 384-well plate and incubated with 10 μM of the indicated compounds for 24 hours. An ANOVA was performed and a Wilcoxon t-test with Holm correction for multiple testing to compare control and compound treated wells. ** = p<0.01. B) Quantification of Wnt activity using the TOPFlash assay in HBEC3-KT cells treated with kenpaullone, 1-azakenpaullone, BIO, or BIO and iCRT14 (a Wnt pathway inhibitor) for 24 hours. Negative control FOPFlash assays performed in parallel were negative as expected (not shown). An ANOVA was performed and a Wilcoxon t-test with Holm correction for multiple testing to compare control and compound treated wells. * = p<0.05. C) qPCR analysis of the Wnt target genes *LEF1* and *AXIN2*. Primary human airway basal cells (n = 4 donors) were treated with compounds for 24 hours. Relative expression was determined by normalising to the reference genes *RPS13* and *GAPDH*. An ANOVA test was performed per target gene and significant Tukey’s HSD values are shown. D) Quantification of primary human airway basal cell proliferation over six days using the CellTiter-Glo assay. Data are normalised to control wells containing the highest concentration of DMSO used in the experimental conditions. Points are the mean value of quadruplicate wells per donor (n = 5 donors). E) Effect of the Wnt pathway inhibitor iCRT14 on the proliferation of airway basal cells in the presence of kenpaullone, 1-azakenpaullone or BIO. Cells were pre-treated with 10 μM iCRT14 or DMSO control for 24 hours before the addition of the Wnt activating compounds. Relative growth was assessed after two days using the CellTiter-Glo assay (n = 4 donors). Wilcoxon tests were performed with Holm correction for multiple testing per compound, ****p<0.0001. F) Quantification of mean organoid size per well with 12 replicate wells per condition were normalised to mean control well organoid size for each donor (n = 5 donors [ID2a = circles, ID5 = triangles, ID6 = squares, ID7 = cross, ID8 = checked box]. An ANOVA was performed per compound and significant Tukey’s HSD values are shown. ** = p<0.01, **** = p<0.0001). Control well data are repeated per compound facet and kenpaullone data is repeated from Figure 2C. G) Immunofluorescence staining showing KRT5 (basal cells, white), MUC5AC (mucosecretory cells, magenta), and ACT (ciliated cells, yellow) in organoids following culture in medium containing the indicated compounds (n=5, donor ID= 8 shown). Scale bars = 50 μm.

### 1-Azakenpaullone activates Wnt signalling in mouse trachea and lung in vivo

Since 1-azakenpaullone had previously been administered *in vivo* in mice (Feng et al., 2012), we performed an initial assessment of the potential of 1-azakenpaullone to modulate airway basal cell Wnt activation and proliferation *in vivo*. We administered 1-azakenpaullone (3 mg/kg) or PBS along with EdU intraperitoneally in C57Bl/6 mice (**Fig. 4A**). 24 hours following administration, we found increased proliferation as judged by EdU positivity within the epithelium (**Fig. 4B**). We also found trends towards increased expression of the Wnt target genes *Myc* and *Ccnd1* in tracheal cells, and a significant increase in *Myc*, but not *Ccnd1* in lung tissue from the same mice (**Fig. 4C**). No changes were observed in *Axin2* or *Lef1* expression in either tissue (**Fig. S4C**). This was supported by increased protein expression of c-Myc and cyclin D1 in 1-azakenpaullone-treated mouse lung tissue (**Fig. 4D**).

**Figure 4:**
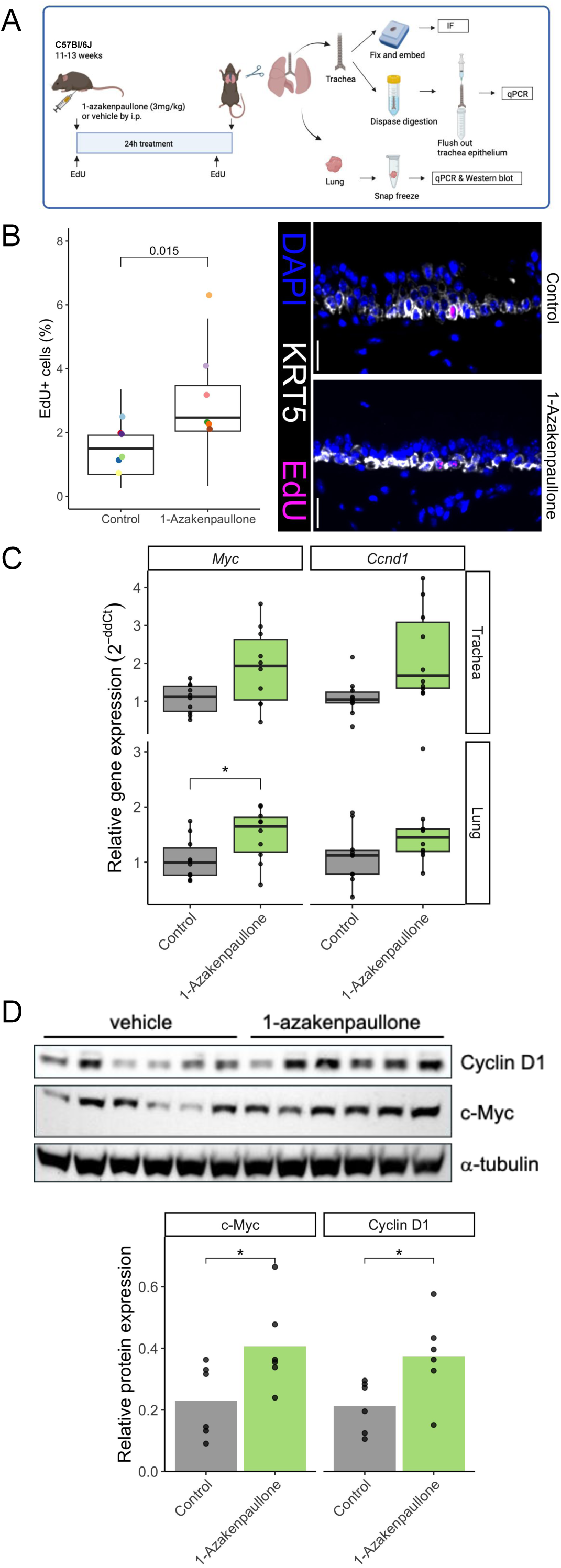
1-Azakenpaullone induced tracheal proliferation and the expression of Wnt target genes *in vivo*. A) Schematic representation of the *in vivo* experimental design. Male C57Bl/6 mice (n = 10 in each group) were administered EdU, and 1-azakenpaullone (3 mg/kg body weight) or vehicle control by intraperitoneal injection. After 24 hours, mice were sacrificed and tracheas and lungs were harvested for analysis. B) Quantification of tracheal proliferation by EdU staining in mice treated with 1-azakenpaullone (3 mg/kg body weight) or vehicle control (n = 6 mice/condition; left). Points indicate mean values per mouse and boxplot represents data from individual images. Wilcoxon test p value shown. Immunofluorescence staining showing KRT5 (basal cells, white) and EdU (magenta) om mouse tracheal sections (right). Scale bars= 20 μm. C) Expression of the Wnt target genes *Myc and Ccnd1* in mouse trachea or and lung as determined by qPCR. Relative expression was normalised to *HPRT1* and/or *ACTB.* A Wilcoxon test was performed (n = 10 mice per group, * p < 0.05). D) Expression of Wnt target proteins in lung tissue as determined by Western blot (n = 6 per group). Alpha-tubulin is shown as a loading control. The bands were quantified using ImageJ and the relative expression of c-Myc and CyclinD1 normalised to alpha-tubulin are shown. A Wilcoxon test was performed, * p < 0.05.

Overall, this study confirms canonical Wnt pathway activation as a pro-proliferative signal in airway epithelial cells cultures and our *in vivo* study demonstrates Wnt activation within the trachea and lung following 1-azakenpaullone administration to uninjured mice. Given that there are 19 Wnt ligands in humans (Rim et al., 2022), Wnt signalling regulation during airway regeneration is likely to be complex, as well as spatiotemporally controlled. The source of Wnt signals *in vivo* seems to differ between airway regeneration (stromal-derived) and homeostasis (airway epithelium-derived) (Aros et al., 2020b). Additionally, non canonical (β-catenin-independent) Wnt signalling can counteract the canonical pathway to inhibit regeneration in alveolar tissue (Baarsma et al., 2017). Future considerations for manipulating the Wnt pathway to improve airway regeneration should account for potential differences in the effect of pathway activation during injury and homeostasis, as previous studies demonstrate that Wnt signalling acts to maintain ΔNp63-expressing stem cells during homeostasis (Haas et al., 2019), with high signalling levels leading to basal cell hyperplasia that is reversible by Wnt inhibition (Aros et al., 2020a). Indeed, constitutive activation of β-catenin during development impacts epithelial cell morphogenesis and leads to polyp formation (Li et al., 2009), persistent Wnt signalling leads to squamous differentiation of airway basal cells (Zhang et al., 2023), and human airway premalignant lesions and squamous cell carcinomas show evidence of Wnt activation (Aros et al., 2020a).

## MATERIALS AND METHODS

### Cell culture

Patient tissue was obtained from regions of histologically normal airway via endobronchial biopsy or as segments of normal lung following lobectomy procedures with informed patient consent. Ethical approval was obtained through the National Research Ethics Committee (REC reference 18/SC/0514). Cell culture plates and flasks were Nunc branded and purchased from Thermo Fisher Scientific. Cells were cultured at 5% CO_2_, 21% O_2_ at 37°C.

3T3-J2 mouse embryonic fibroblasts (a gift from Prof Fiona Watt, King’s College London, U.K.) were cultured in Dulbecco’s Modified Eagle Medium (DMEM; Gibco, 41966) with 9% bovine serum (26170043, Gibco) and 1x penicillin/streptomycin. Feeder layers were generated between passages 9-12. Cells were mitotically inactivated by treatment with 4 μg/mL mitomycin C (M4287, Sigma-Aldrich) in cell culture medium for three hours. Cells were trypsinized and plated at 20,000-30,000 cells/cm^2^. Epithelial cells were added the following day.

Endobronchial biopsies and lobectomy tissue (**Table 1**) arrived in the laboratory on ice in transport medium (αMEM (22561, Gibco) supplemented with 1x penicillin/streptomycin (15140-122, Gibco), 10 μg/mL gentamicin (15710-049, Gibco), and 250 ng/mL amphotericin B (BP264550, Fisher Bioreagents).

**Table 1:**
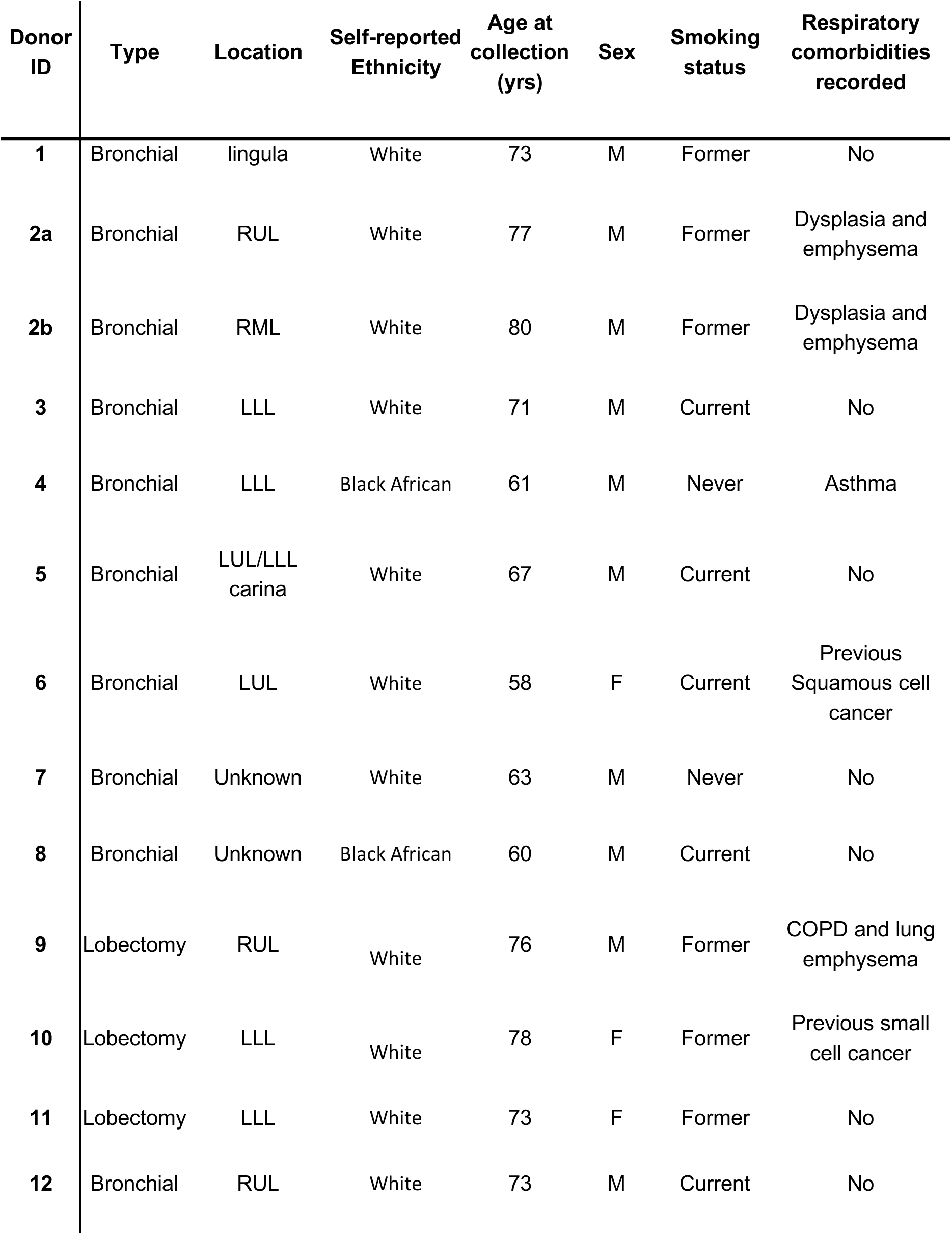
Donor information for primary human airway basal cell cultures.

| Donor ID | Type | Location | Self-reported Ethnicity | Age at collection (yrs) | Sex | Smoking status | Respiratory comorbidities recorded |
| --- | --- | --- | --- | --- | --- | --- | --- |
| 1 | Bronchial | lingula | White | 73 | M | Former | No |
| 2a | Bronchial | RUL | White | 77 | M | Former | Dysplasia and emphysema |
| 2b | Bronchial | RML | White | 80 | M | Former | Dysplasia and emphysema |
| 3 | Bronchial | LLL | White | 71 | M | Current | No |
| 4 | Bronchial | LLL | Black African | 61 | M | Never | Asthma |
| 5 | Bronchial | LUL/LLL carina | White | 67 | M | Current | No |
| 6 | Bronchial | LUL | White | 58 | F | Current | Previous Squamous cell cancer |
| 7 | Bronchial | Unknown | White | 63 | M | Never | No |
| 8 | Bronchial | Unknown | Black African | 60 | M | Current | No |
| 9 | Lobectomy | RUL | White | 76 | M | Former | COPD and lung emphysema |
| 10 | Lobectomy | LLL | White | 78 | F | Former | Previous small cell cancer |
| 11 | Lobectomy | LLL | White | 73 | F | Former | No |
| 12 | Bronchial | RUL | White | 73 | M | Current | No |

To generate single cell suspensions, biopsies and airway tissue were incubated in 16 U dispase (354235, Corning) in RPMI (21875, Gibco) for 20 minutes at room temperature and then mechanically disrupted using sterile forceps. This was followed by incubation in 0.1% trypsin/EDTA (59418C, Sigma-Aldrich) for 30 minutes at 37°C. Both dispase and trypsin/EDTA incubations were neutralised with FBS. The digested tissues were filtered through a 100 μm cell strainer (352360, Falcon). Single cell suspensions were centrifuged at 300 x g for 5 minutes and resuspended in relevant media for counting and plating.

Epithelial cell culture medium (Butler et al., 2016; Hynds et al., 2019; Liu et al., 2012) consisted of a 3:1 ratio of DMEM supplemented with 9% foetal bovine serum (FBS) (10270106, Gibco) and 1x penicillin/streptomycin with Ham’s F12 (21765, Gibco); supplemented with 5 μM Y-27632 (T1725, Tebu-Bio), 25 ng/mL hydrocortisone (H0888, Sigma-Aldrich), 0.125 ng/mL epidermal growth factor (EGF; PHG0315, Gibco), 5 μg/mL insulin (I5500, Sigma-Aldrich), 0.1 nM cholera toxin (C8052, Sigma-Aldrich), 250 ng/mL amphotericin B (BP2645, Fisher Bioreagents), 10 μg/mL gentamicin.

FAD medium consisted of DMEM/Ham’s F12 in a 3:1 ratio supplemented with 10% FBS, 1x penicillin/streptomycin, 1% adenine (A3159, Sigma-Aldrich), 0.5 μg/mL hydrocortisone, 10 ng/mL EGF, 5 μg/mL insulin, 0.1 nM cholera toxin, 2 x 10-5 T3 (T5516, Sigma-Aldrich), 10 μg/mL gentamicin and 250 ng/mL amphotericin B.

The differential trypsin sensitivity of 3T3-J2 fibroblasts and epithelial cells was used to isolate epithelial cell populations from co-cultures. Co-cultures were washed with phosphate-buffered saline (PBS; D8537, Sigma-Aldrich) and 0.05% trypsin/EDTA (25300054, Gibco) was added for two minutes to remove the 3T3-J2 fibroblasts, which are more sensitive to trypsin than the strongly adherent epithelial cells. Flasks were then washed with PBS and 0.05% trypsin/EDTA was added a second time for 5-10 minutes to detach the epithelial cells. Cells were centrifuged at 300 x g for five minutes and resuspended in the required media for counting and plating.

The HBEC3-KT cell line (CRL-4051, ATCC) was cultured in Airway Epithelial Cell Basal Medium (PCS-300-030, ATCC) supplemented with the Bronchial Epithelial Cell Growth Kit (PCS-300-040, ATCC) or in Keratinocyte SFM (17005-042, Thermo Fisher Scientific), a serum-free medium supplemented with recombinant EGF and bovine pituitary extract. Cells were passaged by trypsinization with 0.05% trypsin/EDTA. Cells were centrifuged at 300 x g for five minutes to ensure trypsin removal.

Cells cultured in epithelial cell culture medium were cryopreserved in 50% cell culture medium, 42.5% ProFreeze (BP12-769E, Lonza) and 7.5% DMSO. Cells cultured in FAD medium were frozen in 1x Cell Freezing Medium-Glycerol (C6039, Merck). Cryovials were transferred to a CoolCell and placed in an −80°C freezer to cool by one degree celsius per minute. Cryopreserved cells were then stored at −150°C.

### Virus production

HEK293Ts were cultured in DMEM supplemented with 10% FBS and 1x penicillin/streptomycin. Antibiotics were omitted for one passage prior to and during transfection.

Viral supernatants were produced by co-transfecting HEK293T cells at 70-80% confluency in a T175 flask with 20 μg pHIV-Luc-ZsGreen (gift from Bryan Welm (Addgene plasmid #39196; http://n2t.net/addgene:39196; RRID:Addgene_39196)), 13 μg pCMVR8.74 (gift from Didier Trono (Addgene plasmid #22036; http://n2t.net/addgene:22036; RRID:Addgene_22036)) and 7 μg pMD2.G (gift from Didier Trono (Addgene plasmid #12259; http://n2t.net/addgene:12259; RRID:Addgene_12259)) using JetPEI (101000053, Polyplus Transfection) and following the manufacturer’s protocol. Viral supernatants were collected 48 hours and 72 hours post-transfection and filtered through a 0.45 μm filter (6780-2504, Whatman). Supernatant was combined with PEGit concentrator (5X) (LV810A-1, System Biosciences) overnight at 4°C and centrifuged at 1500 x g for 45 minutes at 4°C. The supernatant was removed, and the viral pellet was resuspended in 1/10th of the original supernatant volume in DMEM supplemented with 25 mM HEPES. Concentrated supernatants were stored at −80°C until use.

### Lentiviral transduction of primary airway basal cells

Epithelial cells were plated at 150,000 cells per well in a 6-well plate on 3T3-J2 feeder layers. The following day, cells were transduced with the addition of 100 μL concentrated virus and polybrene (4 μg/mL; Santa Cruz) to the cell culture medium. Medium was replaced 7 hours after the addition of the virus.

Transduced cells were enriched by fluorescence-activated cell sorting (FACS) using their expression of zsGreen. Single cell suspensions of epithelial cells were collected by differential trypsinization and were filtered through a 70 μm strainer (352350, Falcon), centrifuged 300 x g for five minutes, and resuspended in FACS buffer (PBS, 1% FBS, 25 mM HEPES and 1 mM EDTA). FACS experiments were performed using either a BD FACS Aria or a BD FACS Aria Fusion sorter.

### Compound screening in primary airway basal cells

1,429 compounds from the ENZO library (159 compounds) and the Prestwick Chemical library (1277 compounds) were tested. 7 compounds were shared between both libraries, giving a total of 1,429 unique compounds in our study. Compounds were pre-plated in a 384-well format at 2 mM in DMSO. The plates were stored in a nitrogen-purged StoragePod Roylan Developments) in 5% humidity with 5% O_2_. Potential hit compounds were re-ordered for validation experiments. Compounds (**Table S3**) were reconstituted as 10 mM stocks in DMSO, aliquoted and stored at −20°C to avoid multiple freeze-thaw cycles.

Primary airway basal cells transduced with the pHIV-Luc-ZsGreen construct were expanded in epithelial cell culture medium and cultured for one passage in FAD medium (without Y-27632) prior to plating in 384-well plates (142761, Thermo Fisher Scientific). Compound libraries and DMSO were added to the assay plate using an ECHO 550 liquid handler (Labcyte). Cells were seeded at 2,000 cells/well in FAD medium using a plate dispenser under sterile conditions. After four days of culture, medium was removed, and luciferin was added to cells at 150 μg/mL in FAD medium using the plate dispenser. After ten minutes, bioluminescence was measured using a Envision II plate reader (PerkinElmer). Relative light units were normalised to the DMSO control well values. Data analysis was performed in R. For nuclei visualisation medium was removed and 2 µg/mL Hoechst 33342 (H3570, Invitrogen) in FAD medium was added. After ten minutes the wells were imaged and nuclei counted using a Cytation 3 imaging reader (Biotek).

### Cell proliferation experiments

For experiments in non-transduced HBECs, primary airway basal cells were expanded on 3T3-J2 feeder cells in FAD medium. For proliferation assays, HBECs (without feeder cells) were seeded at 2,000 cells/well in 384-well plates using a E1-ClipTip multichannel pipette (Thermo Fisher Scientific) and compounds were added at the time of seeding. After six days in culture, cells were lysed using CellTiter-Glo (G7570, Promega) reagents, according to the manufacturer’s protocol. Bioluminescence was measured using a Envision II plate reader.

To investigate the effect of inhibiting canonical Wnt activation, primary airway basal cells that had been cultured in FAD medium prior to the assay were seeded at 2,000 cells/well in a 384-well plate in FAD medium containing 10 μM inhibitor of β-catenin responsive transcription 14 (iCRT14; HY-16665, MedChemExpress). The following day hit compounds were added to the wells. After six days in culture, the cell viability was measured using the CellTiter-Glo protocol; the relative light unit was measured by a Envision II plate reader.

### TOPFlash assay

The HBEC3-KT cell line was seeded in 6-well plates at 900,000 cells/well. The following day cells were co-transfected with the M50 Super 8x TOPFlash (a gift from Randall Moon, Addgene plasmid #12456; http://n2t.net/addgene:12456; RRID:Addgene_12456), the M51 Super 8x FOPFlash – an inactive TOPFlash mutant (a gift from Randall Moon, Addgene plasmid #12457) and the pRL SV40 construct (E2231, Promega) using jetOPTIMUS (101000051, Polyplus) according to the manufacturer’s instructions. Medium was changed after 6 hours. The next day, cells were seeded at 20,000 cells/well in a 384-well plate and compounds were added after three hours. Luciferase activity was measured using Dual-Glo luciferase assay system by a Envision II plate reader (PerkinElmer) 48 hours after compound addition.

### Colony formation assays

To calculate colony forming efficiency, primary airway basal cells were seeded at 1,000 cells/well on 6-well plates that had been pre-coated with collagen I and 3T3-J2 feeder cells (Hynds et al., 2018). Experimental compounds or an equivalent DMSO volume for control wells were added the following day. Medium was changed twice per week. On day 10, cells were fixed in 4% PFA for 20 minutes and stained with crystal violet solution (HT90132, Sigma-Aldrich) for 15 minutes at room temperature. Plates were washed thoroughly with water and dried overnight. Plates were scanned using an Epson Perfection V700 PHOTO Scanner. Colonies were manually counted using a brightfield microscope. Colonies were defined as contiguous groups of more than ten cells. Colony forming efficiency (%) was calculated as (number of colonies counted/number of cells seeded) x 100.

### Organoid/bronchosphere culture

Bronchospheres were cultured as per previously published methods (Hynds et al., 2019). Differentiation medium consisted of 50% DMEM and 50% BEBM supplemented with BEGM SingleQuots (except amphotericin B, triiodothyronine, and retinoic acid; CC-3171 and CC-4175, Lonza). Medium was supplemented with 100 nM all-trans retinoic acid (R2625, Sigma-Aldrich) at time of use. Briefly, wells of an ultra-low attachment 96-well plate (3474, Corning) were coated with 30 μL 25% matrigel (354230, Corning) in bronchosphere medium and returned to the incubator at 37°C for 30 minutes. Epithelial cells were seeded at 2,500 cells/well in 65 μL 5% matrigel in bronchosphere medium containing 5 μM Y-27632. Bronchospheres were fed with 50 μL bronchosphere medium supplemented with experimental compounds or equivalent DMSO volume for control wells on days 3, 10 and 17 of culture.

On Day 21, whole well images were taken and image analysis was performed with OrgaQuant (Kassis et al., 2019). Bronchospheres were collected in ice cold PBS and centrifuged at 300 x g for five minutes. Bronchospheres were fixed in 4% PFA on ice for 30 minutes and then centrifuged at 400 x g for five minutes. Bronchospheres were then washed with ice cold PBS and transferred to a well of a V-bottomed 96-well plate. The plate was centrifuged at 400 x g for 5 minutes and bronchospheres were resuspended in 120 µL pre-warmed HistoGel (HG4000012, Fisher Scientific). After ten minutes on ice, the gel was transferred to 70% ethanol at 4°C before being processed in a Leica TP1050 tissue processor. Samples were embedded in type 6 paraffin wax (8336, Epredia) using an embedding station (Sakura Tissue-TEK TEC) and 5 µm sections were cut on a Microm HM 325 microtome.

### Immunohistochemistry, immunofluorescence staining and immunocytochemistry

For immunofluorescence staining of samples on slides, slides were dewaxed using an automated protocol. Slides were washed in PBS and a hydrophobic ring was drawn around the sample using an ImmEdge pen (H-4000, Vector Laboratories). Sections were blocked with 1% bovine serum albumin (BSA; 1.12018.0100, Merck), 5% normal goat serum (NGS; Abcam, ab7481) and 0.1% triton X-100 (X-100, Sigma-Aldrich) in PBS for one hour at room temperature. Primary antibodies raised against TP63 (ab124762, Abcam; 1:300), KRT5 (905901, Biolegend; 1:500), MUC5AC (M5293, Sigma-Aldrich; 1:500) and ACT (T6793, Sigma-Aldrich; 1:500) were diluted in block buffer and applied to slides overnight at 4°C. Slides were washed twice in PBS. For EdU detection the Click-iT^TM^ EdU Cell Proliferation Kit for Imaging, Alexa FluorTM 488 dye (C10337, Thermo Fisher Scientific) was used according to the manufacturer’s protocol prior to secondary antibody application.

Secondary antibodies conjugated to species appropriate Alexa Fluor dyes were diluted 1:1,000 in 5% NGS, 0.1% triton in PBS and applied to slides for three hours at room temperature in the dark. 100 ng/mL DAPI (D9542, Sigma-Aldrich) in PBS was applied to the slides for 20 minutes. Slides were washed twice in PBS and a coverslip was applied manually with Immu-Mount (9990402, Thermo Fisher Scientific). Images were acquired using a Leica DMi8 fluorescence microscope.

For immunocytochemistry, 8-well chamber slides were pre-coated with collagen I (354236, Corning) at 5 μg/cm^2^ in 0.02 N acetic acid (100063, Sigma-Aldrich) for one hour at room temperature. Wells were washed twice with PBS and dried at room temp. Epithelial cells were seeded at 14,000 cells/well. On day 3, cells were fixed in 4% PFA (28908, Thermo Fisher Scientific) for one hour at room temperature and the post-dewax steps of the immunofluorescence protocol above were followed.

### qPCR

Total RNA was extracted using the Quick-RNA Miniprep Plus kit (Zymo Research) according to the manufacturer’s instructions. cDNA was synthesised using qScript cDNA Supermix (95048-100, Quantabio). Quantitative PCR was performed with SYBR Green (4367659, Thermo Fisher Scientific) using QuantStudio5^TM^ Real-time PCR system (Thermo Fisher Scientific). A list of primers used for qPCR is provided in **Table S4**. Statistics were performed on dCt values. Relative gene expression was calculated using a comparative CT method with reference genes (*RPS13* and *GAPDH* for human genes, *Actb and/or Hprt1* for mouse genes).

### In vivo experiment

Animal studies were approved by the University College London Biological Services Ethical Review Committee and licensed under UK Home Office regulations (project licence P36565407). Male mice on the C57Bl/6 background (11-13 weeks of age) were administered one dose of 1-azakenpaullone (3 mg/kg of body weight) or vehicle control (3% DMSO, 50% PEG300, 5% Tween80 and 42% PBS) by intraperitoneal injection. After 24 hours, mice were sacrificed by overdose of pentobarbital for isolation of tracheal and lung tissue. To determine cell proliferation, EdU (Merck) was administered with 1-azakenpaullone 24 hours before harvesting (15 mg/mL of EdU, 50 µg/g body weight), and alone two hours before harvesting (5 mg/ml of EdU, 50 µg/g body weight).

For qPCR analysis of tracheal epithelium, dissected trachea were first incubated in dispase (50 Unit/ml, 354235, Corning) for 40 minutes at 37°C. Following incubation, the epithelial layer was flushed from the trachea with 10 mL ice-cold PBS into a 15 mL tube using a needle and syringe. Cells were centrifuged at 400 x g and resuspended in RNA lysis buffer for RNA extraction.

To enable immunofluorescence staining of the tracheal epithelium, dissected tracheas were fixed in 4% PFA at 4°C overnight and then stored in 70% ethanol at 4°C before being processed in a Leica TP1050 tissue processor. Samples were embedded in type 6 paraffin wax (8336, Epredia) using an embedding station (Sakura Tissue-TEK TEC) and 5 µm sections were cut on a Microm HM 325 microtome.

To automate quantification analyses, macros were created in Fiji (Schindelin et al., 2012) to mask and quantify the count and area of immunofluorescent staining. Counts were performed on tracheal sections by two independent researchers blinded to the condition groups. The mean EdU proportion was taken per section.

### Western blotting

Mouse lungs were snap frozen on dry ice, minced with a scalpel, homogenised and lysed in RIPA lysis buffer with protease inhibitors (cOmplete Ultra tablets, 5892970001, Merck) on ice to extract protein. 30 μg of protein samples were separated by SDS-PAGE and transferred onto nitrocellulose membranes using an iBlot2 Dry Blotting System (Thermo Fisher Scientific). Membranes were incubated with anti-cyclinD1 (55506, Cell Signaling Technologies), anti-c-myc (ab32072, Abcam), or anti-alpha-tubulin (9099, Cell Signaling Technologies) antibodies, washed, incubated with species-appropriate horseradish peroxidase (HRP)-conjugated secondary antibodies. After incubating with the substrate (Immobilon Crescendo Western HRP Substrate; WBLUR0500, Millipore), chemiluminescent signals were visualised using an iBright FL1500 imaging system (Thermo Fisher Scientific). Quantification of bands was performed using ImageJ (Image Processing and Analysis in Java, The National Institute of Health).

## AUTHOR CONTRIBUTIONS

Conceptualization: S.M.J., R.E.H.

Methodology: Y.I., J.C.O., D.R.B., D.R.P., K.A.L., R.E.G., R.K., N.O.C., R.E.H.

Investigation: Y.I., J.C.O., M-B.E.M., D.B.

Resources: R.K., N.O.C., S.M.J.

Writing – original draft preparation: Y.I., J.C.O., R.E.H.

Writing – review and editing: M-B.E.M., D.R.P., N.O.C., S.M.J.

Visualisation: Y.I., J.C.O.

Supervision: S.M.J., R.E.H.

Project administration: R.E.H.

Funding acquisition: M.Z.N., S.M.J., R.E.H.

## ACKNOWLEDGEMENTS

The authors thank Bernadette Carroll (University College London Hospitals, London, U.K.), Dr. Matthew Wright (University College London Hospitals, London, U.K.) and members of the ASCENT study team (UCL Respiratory, University College London, U.K.) for their roles in collecting patient tissue for our study. The authors acknowledge the assistance of Mr Jamie Evans (Division of Medicine, University College London, U.K.) and Mr George Morrow, (UCL Cancer Institute, University College London, U.K.) in performing fluorescence-activated cell sorting experiments.

## DATA AVAILABILITY

All relevant data can be found within the article and its supplemental information.

