## Supplementary Information for "Compound screening in primary human airway basal cells identifies Wnt pathway activators as potential pro-regenerative therapies"

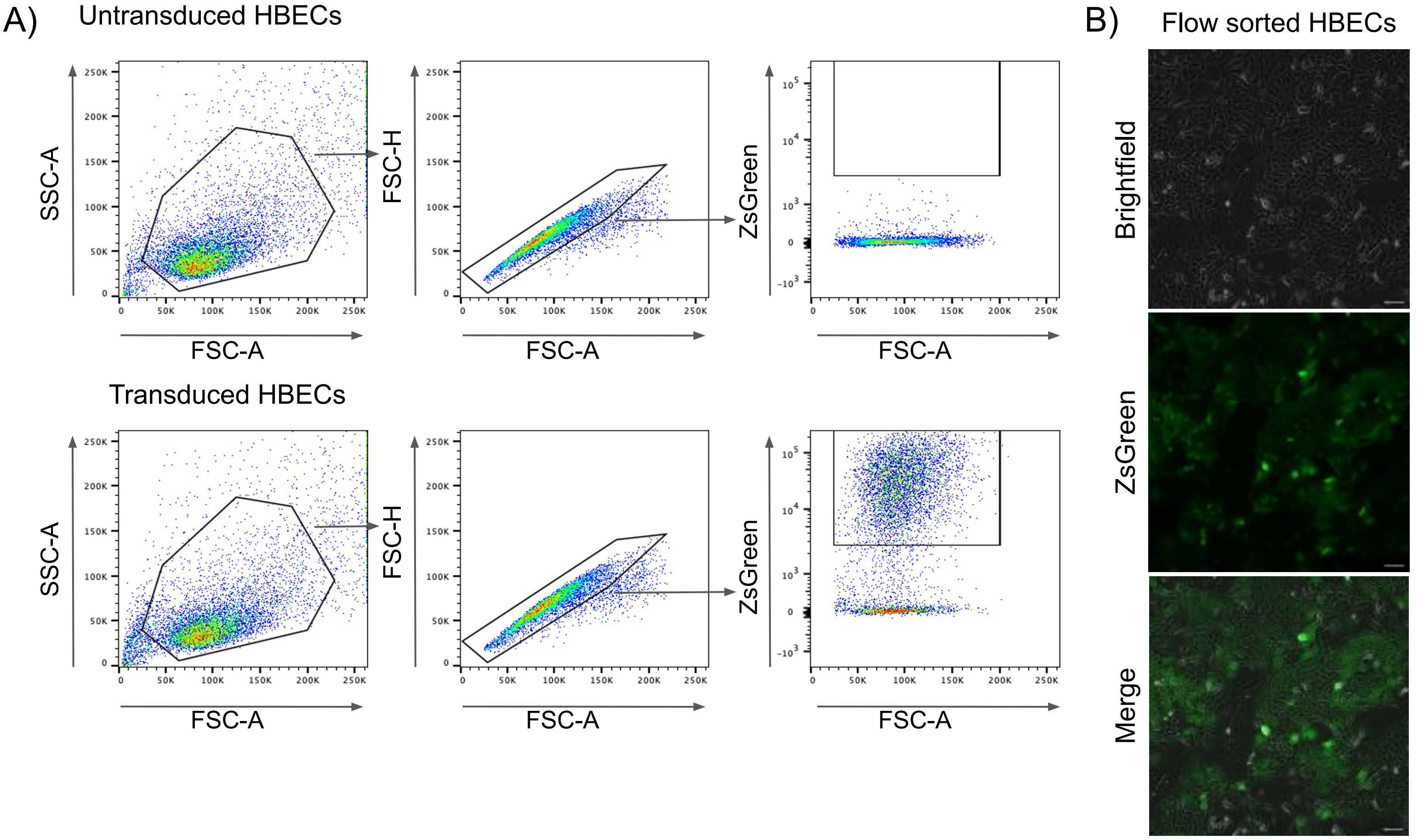


C)

| Donor ID | Transduction efficiency (%) (ZsGreen+ of single cell population) | Number of cells sorted |
|---|---|---|
| 1 | 31 | 270,000 |
| 2a | 65.5 | 200,000 |
| 3 | 68.1 | 100,000 |
| 4 | 72.1 | 75,000 |

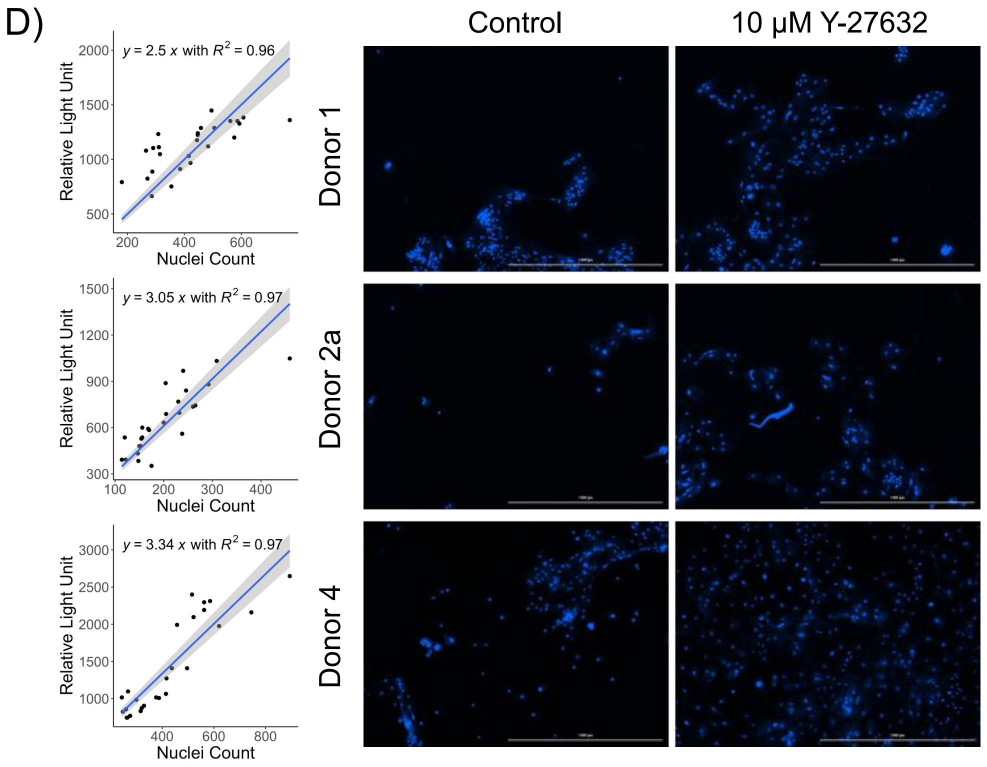


**Figure S1: Transduction and validation of zsGreen-Luciferase transduced primary airway basal cells for compound screening.**

A) Gating strategy for flow sorting of ZsGreen+ population with data shown from donor 4.

B) Representative brightfield and fluorescence images of flow sorted human bronchial epithelial cells (HBECs) from a single donor. Scale bars = 50 μm.

C) Transduction efficiency for each HBEC donor following lentiviral transduction with the pHIV-Luc-Zsgreen construct.

D) Correlation between bioluminescence and nuclei count (n=3 HBEC donors) (left). Each data point is a reading from one well. Representative images of Hoechst 33342 staining of cells following the luciferase assay (right). Scale bars = 1,000 μm.

A)

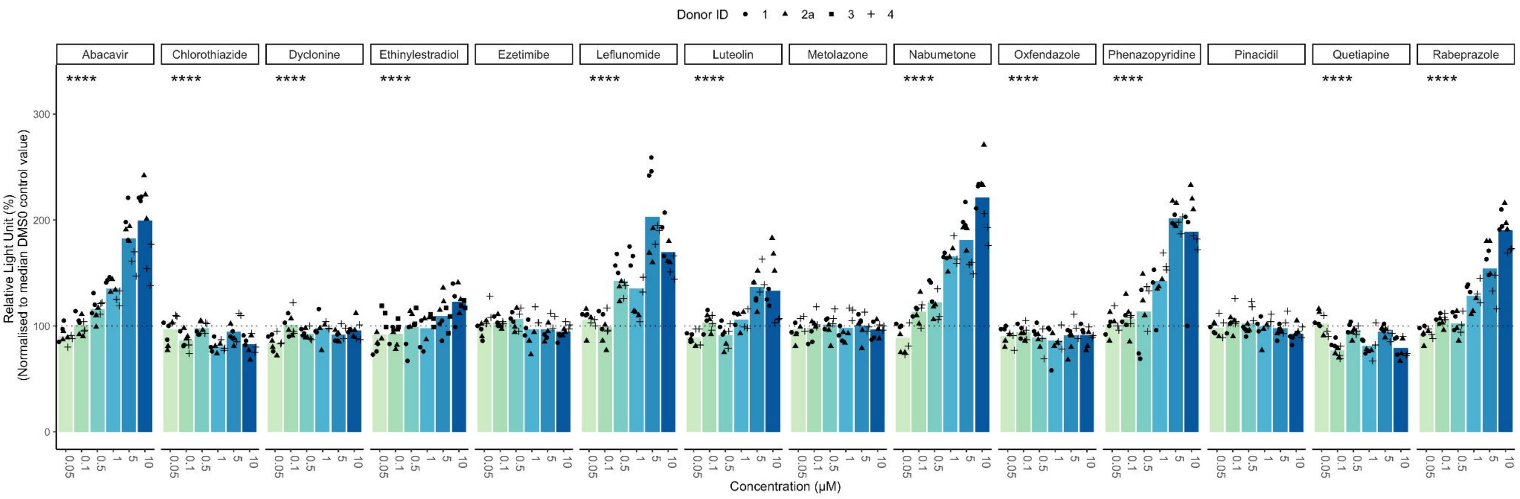


B)

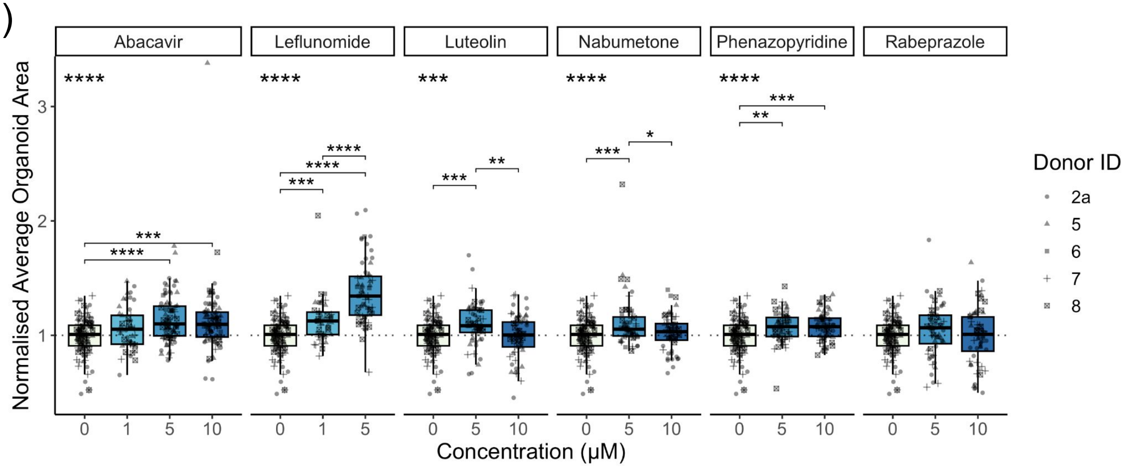


**Supplemental Figure 2: Related to Figure 2.**

A) Four-day concentration-response proliferation assays in primary human airway basal cells transduced with the pHIV-Luc-zsGreen construct for compounds identified within the Prestwick Chemical library (n = 4/5 donors per compound; donor ID1 = circles, ID2a = triangles, ID3 = squares, ID4 = cross). An ANOVA was performed per compound, **** = p<0.0001.

B) Quantification of mean organoid size per well with 12 replicate wells per condition were normalised to mean control well organoid size for each donor (n = 5 donors [donor ID2a = circles, ID5 = triangles, ID6 = squares, ID7 = cross, ID8 = checked box]. An ANOVA was performed per compound and significant Tukey's HSD values are shown. * = p<0.05, ** = p<0.01, ***= p<0.001, **** = p<0.0001). Control well data are repeated per compound facet, data for Nabumetone and Phenazopyridine are repeated in 2C.

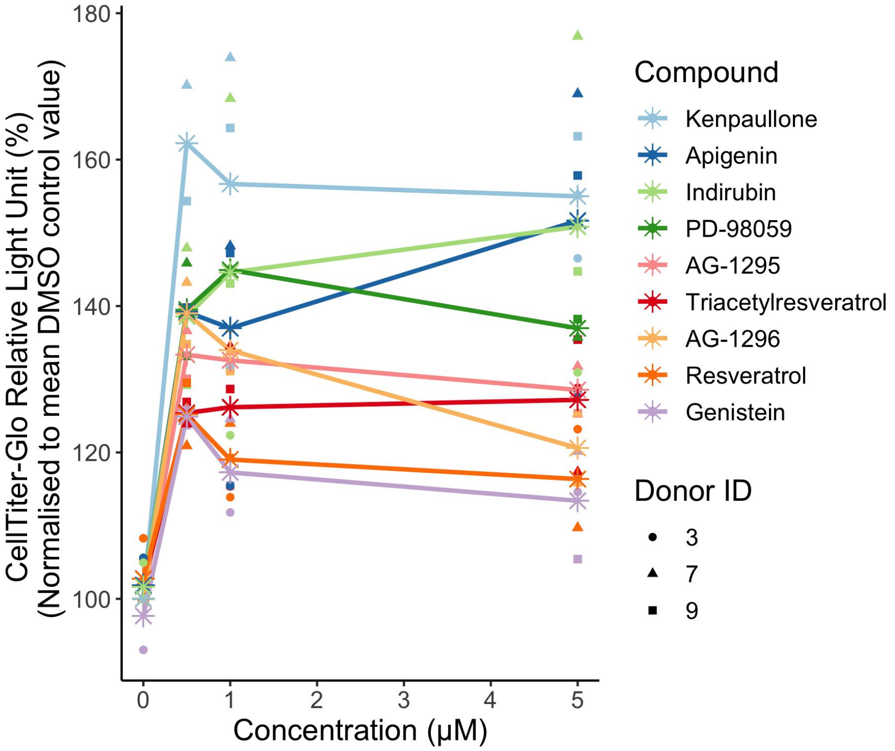


**Supplemental Figure 3: A concentration-response experiment on hit compounds from the ENZO chemical library.** Untransduced primary human bronchial epithelial cells (n=3 donors) were cultured with the indicated concentrations of screen hit compounds for 6 days. Relative cell growth was assessed using the CellTitre-Glo assay. * denotes mean value at each concentration tested.

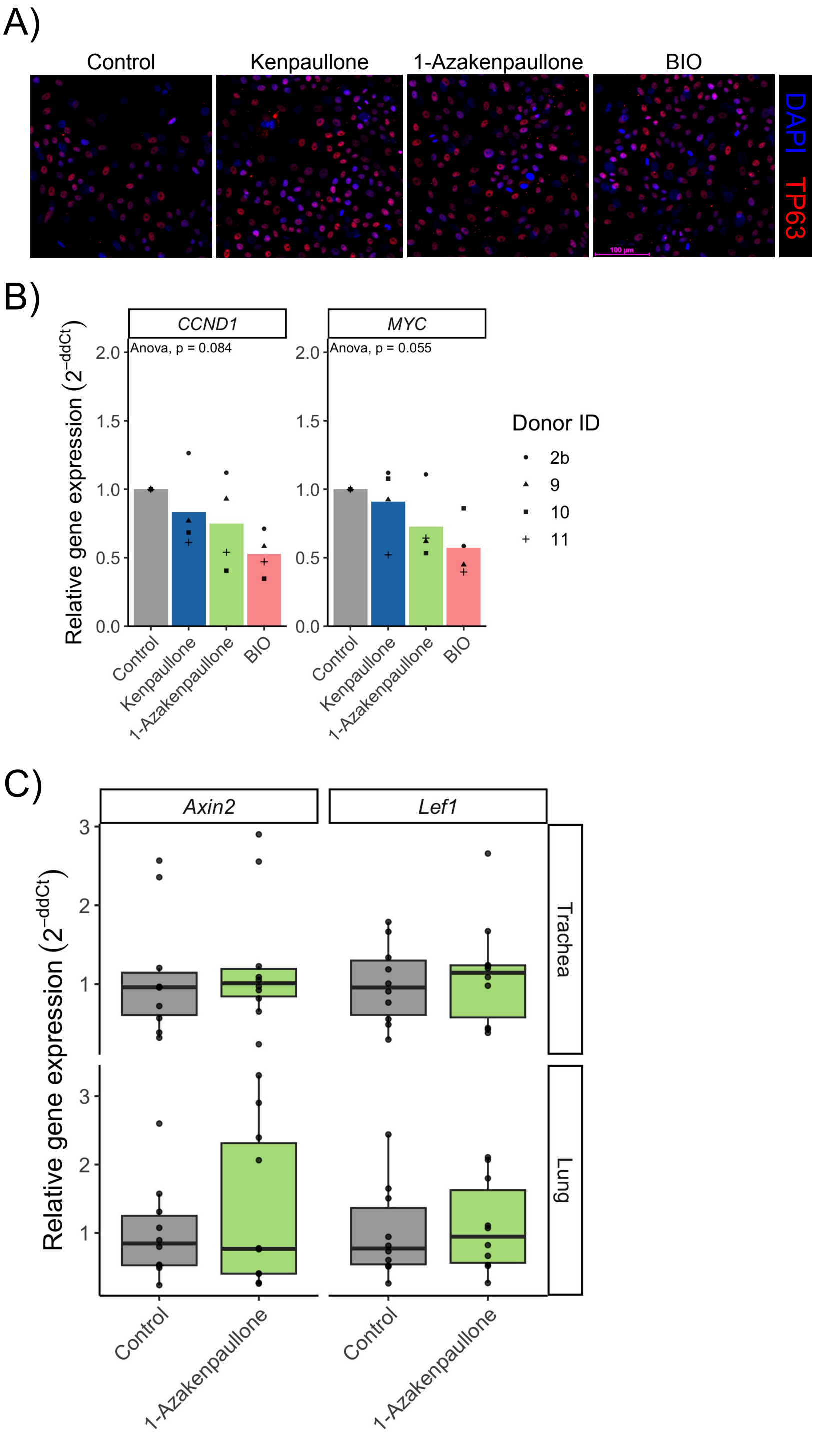


**Supplemental Figure 4: Additional data relating to Figure 3 and Figure 4.**

A) Immunofluorescence staining for TP63 (red) and DAPI (blue) in airway basal cells treated with kenpaullone (5 µM), 1-azakenpaullone (5 µM) or BIO (2 µM) for 24 hours. Scale bar = 100 µm.

B) qPCR analysis of the Wnt target genes *CCND1 and MYC* in primary human airway basal cells (n = 4 donors) treated with compounds for 24 hours. Relative expression was determined by normalising to the reference genes *RPS13* and *GAPDH*. An ANOVA test was performed per target gene.

C) Expression of the Wnt target genes *Axin2 and Lef1* in mouse trachea or lung as determined by qPCR. Relative expression was normalised to *ACTB.* A Wilcoxon test was performed, no significance found (n = 10 mice per group)

| **Compound name** | **Library** | **Compound description/Therapeutic effect (target and target mechanism)** |
|---|---|---|
| Triacetylresveratrol | ENZO; Epigenetics | SIRT1 activator |
| Kenpaullone | ENZO; Kinase Inhibitor | GSK-3β inhibitor |
| AG-1295 | ENZO; Kinase Inhibitor | Tyrosine Kinase inhibitor |
| Y-27632.2HCl | ENZO; Kinase Inhibitor | Rho Kinase inhibitor |
| Resveratrol | ENZO; Epigenetics | SIRT1 activator |
| Apigenin | ENZO; Kinase Inhibitor | CK-II inhibitor |
| PD-98059 | ENZO; Kinase Inhibitor | MEK inhibitor |
| GF 109203X | ENZO; Kinase Inhibitor | PKC inhibitor |
| PP1 | ENZO; Kinase Inhibitor | Src inhibitor |
| AG-1296 | ENZO; Kinase Inhibitor | PDGFRK |
| Indirubin-3'-monoxime | ENZO; Kinase Inhibitor | GSK-3β inhibitor |
| Genistein | ENZO; Kinase Inhibitor | Tyrosine Kinase inhibitor |
| Dyclonine hydrochloride | Prestwick Chemical | Local anesthetic<br>(Voltage-gated NA+ channel inhibitor, Aldehyde dehydrogenase inhibitor) |
| Chlorothiazide | Prestwick Chemical | Antihypertensive, Diuretic<br>(Thiazide-sensitive sodium-chloride cotransporter inhibitor, Carbonic anhydrase I, II, IV inhibitor) |
| Pinacidil | Prestwick Chemical | Antihypertensive, Vasodilator, Anti-inflammatory<br>(K+ channel Ca2+ dependant activator) |
| Leflunomide | Prestwick Chemical | Immunosuppressant, Antineoplastic<br>(Dihydroorotate dehydrogenase inhibitor) |
| Rabeprazole sodium salt | Prestwick Chemical | Antiulcer<br>(H+/K+ ATPase) |
| Ceftazidime pentahydrate | Prestwick Chemical | Antibacterial<br>(Penicillin-binding protein 1A, 1B, 2,3,4 inhibitor) |
| Phenazopyridine hydrochloride | Prestwick Chemical | Analgesic, Antidotes<br>(microtubule associated protein tau inhibitor, estrogen receptor 1 agonist, nuclear receptor 1H4 antagonist) |
| Abacavir Sulfate | Prestwick Chemical | Antiviral<br>(Reverse transcriptase inhibitor) |
| Nabumetone | Prestwick Chemical | Analgesic, Anti-inflammatory<br>(Cyclooxygenase) |
| Metolazone | Prestwick Chemical | Antihypertensive, Diuretic, Antineoplastic<br>(Thiazide-sensitive sodium-chloride cotransporter inhibitor) |
| Quetiapine hemifumarate | Prestwick Chemical | Antipsychotic<br>(Dopaminergic and 5-HT receptors antagonist) |
| Oxfendazole | Prestwick Chemical | Antihelmitic |
| Ethinylestradiol | Prestwick Chemical | Contraceptive<br>(Estrogen receptor) |
| Luteolin | Prestwick Chemical | Expectorant, Antineoplastic |
| Ezetimibe | Prestwick Chemical | Hypocholesterolemic<br>(Niemann-Pick C1-like protein 1) |

**Table S1: Compound descriptions for 27 hit compounds.**

Ordered as Fig. 1E

Table S2: Z Scores for all screened compounds.

Data found in Screen_Results.xlsx.

| Compound | Supplier(s) | Catalogue number |
|---|---|---|
| Resveratrol | ENZO | BML-FR104-0100 |
| Triacetylresveratrol | ENZO | BML-FR119-0010 |
| Apigenin | ENZO | BML-EI345-0020 |
| Kenpaullone | ENZO<br>Cayman Chemical | BML-EI310-0005<br>10010239 |
| Indirubin-3'-monoxime | ENZO | BML-CC207-0001 |
| PD-98059 | ENZO | BML-EI360-0005 |
| AG-1296 | ENZO | BML-EI303-0005 |
| GF 109203X | ENZO | BML-EI246-0001 |
| TYRPHOSTIN AG 1295 | ENZO | ALX-270-035-M001 |
| Genistein | ENZO | ALX-350-006-M010 |
| Hypericin | ENZO | ALX-350-030-M001 |
| Phenazopyridine (hydrochloride) | Cayman Chemical | 29683 |
| Nabumetone | Cayman Chemical | 20251 |
| Rabeprazole (sodium salt) | Cayman Chemical | 14939 |
| Dyclonine (hydrochloride) | Cayman Chemical | 27667 |
| Luteolin | Cayman Chemical | 10004161 |
| Metolazone | Cayman Chemical | 15987 |
| Leflunomide | Cayman Chemical | 14860 |
| Quetiapine (hemifumarate) | Cayman Chemical | 14152 |
| Chlorothiazide | Cayman Chemical | 17909 |
| Dichlorphenamide | Cayman Chemical | 23658 |
| Oxfendazole | LKT Laboratories | 09322 |
| Ezetimibe | Adooq Bioscience | A10379 |
| Pinacidil monohydrate | Santa Cruz Biotechnology | SC-203198 |
| BIO | Cayman Chemical | 13123 |
| 1-Azakenpaullone | Cayman Chemical | 16733 |

**Table S3: Compounds used for validation of screening results.**

| Gene | Species | Primer sequence (5'-3') |
|---|---|---|
| *LEF1* | human | Forward: AGAACACCCCGATGACGGA<br>Reverse: GGCATCATTATGTACCCGGAAT |
| *AXIN2* | human | Forward: CAACACCAGGCGGAACGAA<br>Reverse: GCCCAATAAGGAGTGTAAGGACT |
| *CCND1* | human | Forward: CCGAGAAGCTGTGCATCTACAC<br>Reverse: AGGTTCCACTTGAGCTTGTTCAC |
| *MYC* | human | Forward: CCTGGTGCTCCATGAGGAGAC<br>Reverse: CAGACTCTGACCTTTTGCCAGG |
| *RPS13* | human | Forward: TCGGCTTTACCCTATCGACGCAG<br>Reverse: ACGTACTTGTGCAACACCATGTGA |
| *GAPDH* | human | Forward: TGATGACATCAAGAAGGTGGTGAAG<br>Reverse: TCCTTGGAGGCCATGTAGGCCAT |
| *Lef1* | mouse | Forward: TGTTTATCCCATCACGGGTGG<br>Reverse: CATGGAAGTGTCGCCTGACAG |
| *Axin2* | mouse | Forward: TGACTCTCCTTCCAGATCCCA<br>Reverse: TGCCCACACTAGGCTGACA |
| *Ccnd1* | mouse | Forward: GCAGAAGGAGATTGTGCCATCC<br>Reverse: AGGAAGCGGTCCAGGTAGTTCA |
| *Myc* | mouse | Forward: CAGAGGAGGAACGAGCTGAAGCGC<br>Reverse: TTATGCACCAGAGTTTCGAAGCTGTTCG |
| *Actb* | mouse | Forward: ATGGCTGGGGTGTTGAAGGT<br>Reverse: ATCTGGCACCACACCTTCTACAA |
| *Hprt1* | mouse | Forward: CTGGTGAAAAGGACCTCTCGAAG<br>Reverse: CCAGTTTCACTAATGACACAAACG |

**Table S4: qPCR primer sequences.**
